# Partial epithelial-to-mesenchymal transition is prognostic and associates with Slug in head and neck cancer

**DOI:** 10.1101/2020.10.20.346692

**Authors:** Henrik Schinke, Min Pan, Merve Akyol, Jiefu Zhou, Gisela Kranz, Darko Libl, Florian Simon, Martin Canis, Philipp Baumeister, Olivier Gires

## Abstract

Therapy resistance leading to local recurrence and metastases remains highly problematic in head and neck squamous cell carcinomas (HNSCC). Single cell RNA-sequencing defined a partial epithelial-to-mesenchymal transition (p-EMT) signature associated with metastases in HNSCC. However, the prognostic value of the p-EMT signature and potential drivers of p-EMT in HNSCC remain unclear. Here, single sample scoring of molecular phenotypes (Singscoring) served to establish clinical p-EMT-Singscores that were significantly associated with nodal metastases and predicted overall survival in two independent HNSCC cohorts. p-EMT-Singscores correlated most strongly with EMT transcription factor (EMT-TF) Slug. *In vitro*, Slug promoted p-EMT, enhanced invasion, and resistance to irradiation. In patients, Slug protein levels in tumors predicted disease-free survival and its peripheral expression at the interphase to tumor-microenvironment was significantly increased in recurring patients. Thus, p-EMT represents a novel clinical risk-predictor that impacts on HNSCC patients’ outcome and is partly controlled by Slug.

## Introduction

Every year over 600,000 patients are diagnosed with head and neck squamous cell carcinoma (HNSCC) (Ferlay et al., 2015; Siegel, Miller, & Jemal, 2016). HNSCC are challenging to treat due to their aggressive features in combination with the high functional and aesthetic relevance of the head and neck region that can limit radical therapeutic approaches. HNSCC show high levels of phenotypic and molecular inter- and intratumor heterogeneity (Cancer Genome Atlas, 2015; Chung et al., 2004; Leemans, Snijders, & Brakenhoff, 2018; Puram et al., 2017; Stransky et al., 2011; Walter et al., 2013) that is associated with poor prognosis and five-year OS rates below 50 % (Mroz, Tward, Hammon, Ren, & Rocco, 2015; Mroz et al., 2013). In addition to a separation of human papillomavirus (HPV)-positive and -negative HNSCC, gene expression and mutational analyses resulted in the definition of classical, atypical, basal-like, and mesenchymal molecular subtypes of HNSCC (Cancer Genome Atlas, 2015; Keck et al., 2015; Stransky et al., 2011; Walter et al., 2013).

A major cellular differentiation program responsible for heterogeneity in several tumor entities is the epithelial-to-mesenchymal transition (EMT) (Brabletz, Kalluri, Nieto, & Weinberg, 2018; Guo et al., 2014; Lambert, Pattabiraman, & Weinberg, 2017). In the process, apical-basal polarity of epithelial cells with tight cell-to-cell junctions is gradually lost and cells adopt a spindle-like morphology associated with elevated motility and invasiveness (Lim & Thiery, 2012; Nieto, Huang, Jackson, & Thiery, 2016). Mesenchymal features support tumor cells to complete various steps of the metastatic cascade, including local invasion, intravasation into blood and lymphatic vessels, re-localization within the patient’s body *via* circulation, extravasation into parenchyma in locoregional and distant tissue, and survival as micro-metastatic deposits (Lambert et al., 2017; Thiery, 2002). Accordingly, growing evidence shows that EMT traits are implicated in metastasis formation and therapy resistance *in vivo*, resulting in poor prognosis for patients (Brabletz et al., 2018; Chaffer, San Juan, Lim, & Weinberg, 2016; Lambert et al., 2017). Controversially, it was observed that EMT might be dispensable for invasion and metastasis but is key for the cells’ resistance to chemotherapeutic drugs (Fischer et al., 2015; Zheng et al., 2015). The observed dispensability of selected EMT transcription factors (EMT-TFs) for metastasis formation might partly be explained by a high complexity and redundancy of regulatory molecular mechanisms along with the temporal resolution and plasticity of the EMT program in cancer (Aiello et al., 2017; Brabletz et al., 2018; Ye et al., 2017). In fact, co-existing intermediate EMT states were found in primary and systemic tumor cells at different phases of the metastatic cascade and at various stages of disease progression and therapy response (Chaffer et al., 2016; Diepenbruck & Christofori, 2016; Lambert et al., 2017; Liu et al., 2019; Nieto & Cano, 2012; Nieto et al., 2016; Yu et al., 2013). Owing to the frequently incomplete, transitional, and reversible nature of EMT in cancer, the term partial EMT (p-EMT) was coined to describe EMT in malignant cells more accurately (Aiello & Kang, 2019; Zhang & Weinberg, 2018).

scRNAseq of a small cohort of oral cavity tumors confirmed a high degree of intratumor heterogeneity and molecularly defined tumor cells in a state of p-EMT (Puram et al., 2017). Amongst a gene set of one hundred genes, a more confined common p-EMT signature of 15 genes was identified in oral cavity tumors of the basal-like subtype, which correlated with the presence of nodal metastases, higher grades, and other adverse pathological parameters (Puram et al., 2017). However, an implementation of the common p-EMT gene signature as a stratifying parameter to predict clinical outcome of HNSCC is pending.

Molecularly, the EMT program can be initiated by the master regulators SNAI1 (SNAIL), SNAI2 (SLUG), TWIST, ZEB1, and ZEB2 that act as transcriptional repressors to regulate genes related to cell adhesion, migration, and invasion (Bolos et al., 2003; Lamouille, Xu, & Derynck, 2014; Peinado, Olmeda, & Cano, 2007). However, regulation of p-EMT in HNSCC remains insufficiently studied. Hence, the present study assessed a potential implementation of p-EMT as a prognostic parameter and further focused on p-EMT regulators in HNSCC.

## Results

### The p-EMT common signature is prognostic in HNSCC and correlates with nodal metastases

The p-EMT gene signature originating from scRNAseq of oral cavity cancers (n = 18) (Puram et al., 2017) has to date not been transferred to bulk sequencing results of independent HNSCC cohorts to prognosticate clinical endpoints. In order to make best use of the clinical feasibility of bulk sequencing and the information content of scRNAseq, HPV-negative patients of the TCGA cohort (n = 243) (Cancer Genome Atlas, 2015) were subdivided into atypical, classical, basal, and mesenchymal molecular subtypes. In accordance with the reported importance of p-EMT in the malignant-basal subtype, as defined by Puram *et al.* (Puram, Parikh, & Tirosh, 2018; Puram et al., 2017), basal (n =38) and mesenchymal subtypes (n = 46), representing patients with high intra-subgroup similarity, were included for further analysis (**Fig. 1A, B**). The influence of non-malignant cells was inferred from the heatmap with hierarchical clustering of a distance matrix of selected marker genes (Puram et al., 2017). Patients in a cluster with highest expression of marker genes of non-malignant cells were excluded as illustrated. The resulting n = 55 patients represented a malignant-basal subtype surrogate within the TCGA cohort (**Fig. 1C**). Expression levels of genes of the common p-EMT signature (n = 15) were visualized in malignant-basal patients, showing a heterogeneous repartition of formerly basal and mesenchymal subtypes (**Fig. 1D**). Of note, a general stratification purely based on raw gene expression values did not allow a prediction of clinical endpoints (data not shown). Therefore, p-EMT-Singscores were generated (Foroutan et al., 2018), based on the relative expression of n = 15 common p-EMT genes against the background of n = 10,000 genes expressed in the malignant basal subtype surrogate in TCGA (**Fig. 1E**). p-EMT-Singscores represent patient-specific, single sample gene set enrichments with values ranging from −1 to 1 defining the degree of p-EMT of individual patients within the TCGA cohort. p-EMT-Singscores resembled a normal distribution as compared to a theoretical quantile distribution (**Suppl. Fig. 1A**, upper left panel). Plotting of the normalized ranks of the n = 15 common p-EMT genes displayed a more widespread density for the patient with the lowest p-EMT Singscore, whereas it showed a stronger accumulation of the genes towards higher ranks in the patient with the highest p-EMT Singscore (**Suppl. Fig. 1A**, lower panels). This accumulation was further substantiated in a dispersion plot, where high p-EMT-Singscores correlated with low dispersion of normalized ranks (**Suppl. Fig. 1A**, upper right panel).

**Figure 1:**
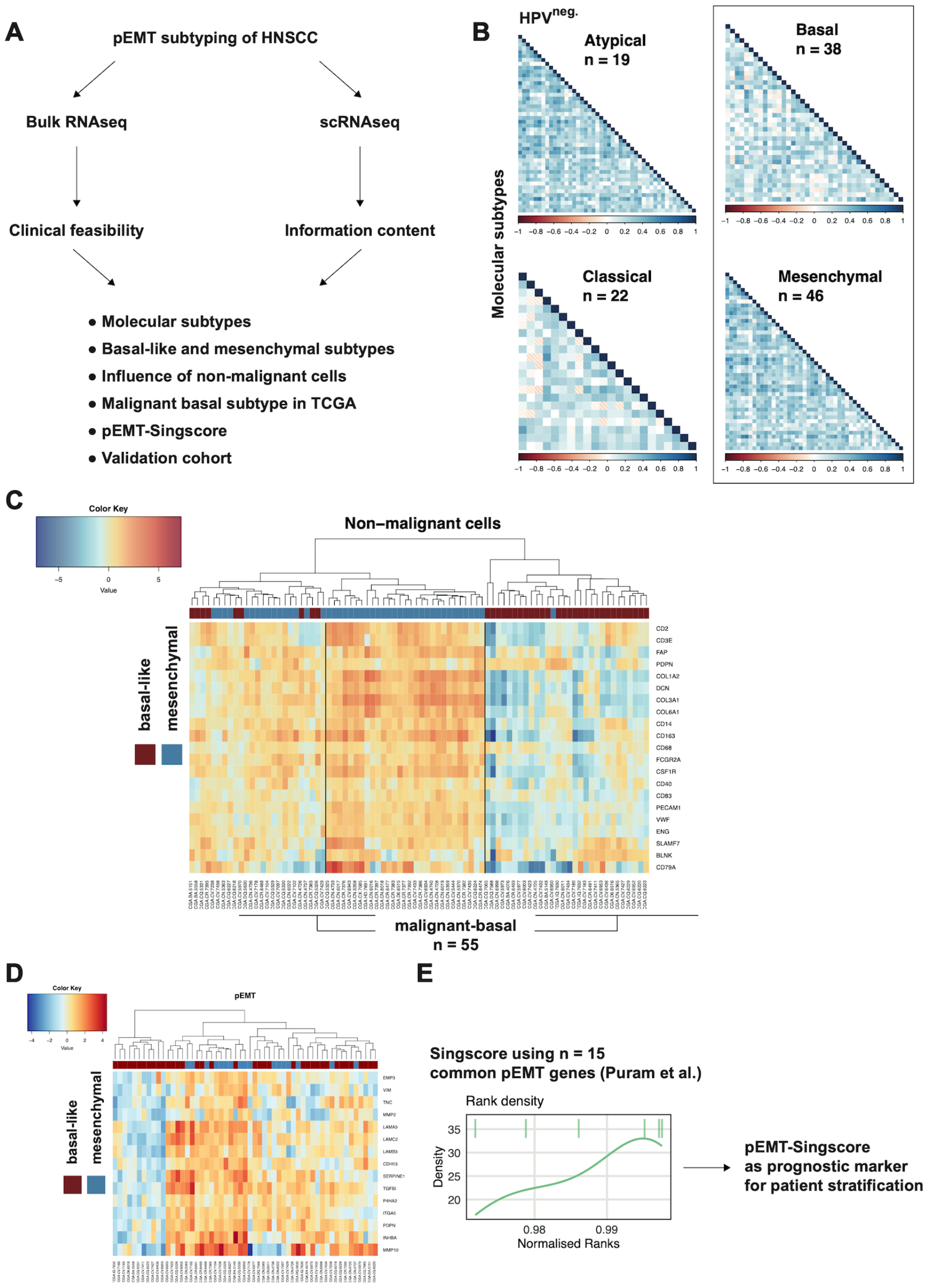
Transfer of scRNAseq p-EMT signature to bulk sequencing using Singscores. (**A**) Schematic representation of p-EMT subtyping of HNSCC using scRNAseq results in the malignant-basal subtype of HNSCC in TCGA. (**B**) Subtype classification of HPV-negative HNSCC of the TCGA cohort into atypical, classical, basal, and mesenchymal. (**C**) Heatmap with hierarchical clustering of Euclidean distance matrix of the expression of marker genes for non-malignant cells within all patients of the basal and mesenchymal subtypes. Patients with a high expression of non-malignant genes were excluded to generate a malignant-basal surrogate cohort (n = 55). Basal and mesenchymal subtypes are marked in blue and red boxes as indicated. (**D**) Heatmap with hierarchical clustering of Euclidean distance matrix of the n = 15 common p-EMT genes (Puram et al., 2017) in the malignant-basal surrogate TCGA cohort. Basal and mesenchymal subtypes are marked in blue and red boxes as indicated. (**E**) Representative examples of a density plot of normalized ranks for genes of the common p-EMT signature used to form p-EMT-Singscores.

Overall survival (OS) after five years of patients within the 1^st^ quartile of p-EMT-Singscores in a Cox proportional hazard model visualized in Kaplan Meier curves was significantly increased compared to patients with medium and high p-EMT-Singscores (2^nd^ and 3^rd^ quartiles, and 4^th^ quartile) (**Fig. 2A**, High vs. low: HR: 6.64, CI: 1.78 – 24.78, logrank p = 0.005; High vs. medium: HR: 2.73, CI: 1.15 – 6.52, logrank p = 0.04). p-EMT-Singscores showed a normal distribution (**Suppl. Fig. 1A, upper left pannel**). The univariate Cox proportional hazard model using p-EMT-Singscores as a feature resulted in a significant model (HR: 785.7, 95% CI: 1.34 – 461665, logrank p = 0.04). To test the specificity of p-EMT-Singscores, calculations were repeated 10,000 times with different 15 randomly selected genes from the extracted TCGA gene pool (n= 10,000 genes excluding all n = 100 p-EMT genes). Plotting of logrank p-values *versus* Cox proportional hazard ratios of all 10,000 random sets of n = 15 genes yielded a total of 4.18 % sets reaching a HR above 1 with a significant p-value of 0.05 or lower (**Fig. 2B**), demonstrating an acceptable alpha error level below 0.05 and the high specificity of the p-EMT-Singscore.

**Fig. 2:**
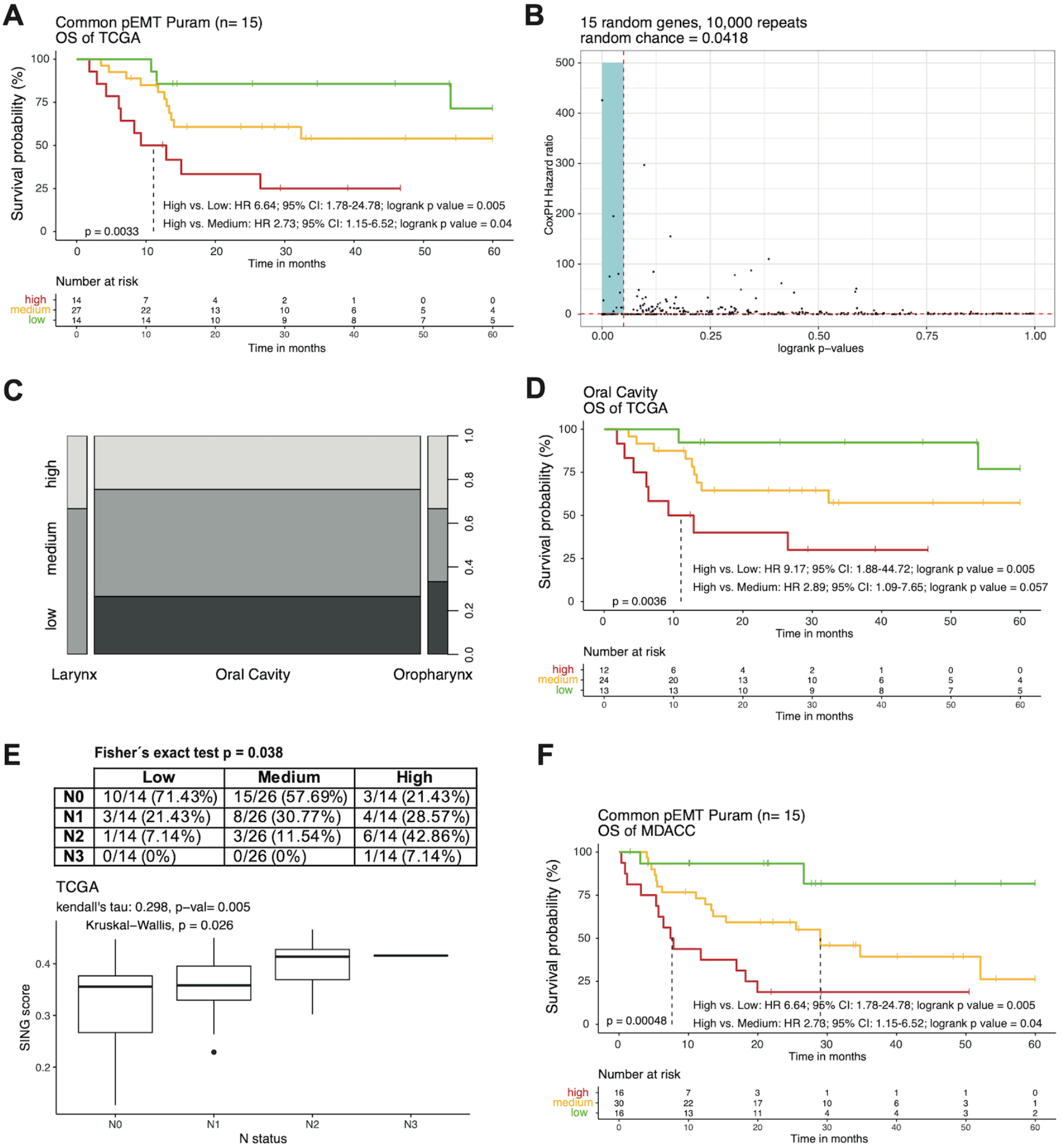
Partial EMT is prognostic in HNSCC. (**A**) Singscores were calculated for n = 15 common p-EMT genes for each of the n = 55 TCGA malignant-basal TCGA patients and served to compute a Cox proportional hazard model. Kaplan-Meier curve stratified after 1^st^ quartile (low), 2^nd^ and 3^rd^ quartiles (medium), and 4^th^ quartile (high) Singscores with Cox proportional hazard ratio (HR), 95% confidence intervals (CI), logrank p-values and survival table is shown. (**B**) Cox proportional hazard models were computed n = 10,000 times with n = 15 random genes (except n = 100 p-EMT genes). Logrank p-values are plotted against HR. Cutoffs indicated logrank p = 0.05 and HR = 1. (**C**) Shown are the repartition of Singscores into 1^st^ quartile (low), 2^nd^ and 3^rd^ quartiles (medium), and 4^th^ quartile (high), and the HNSCC sub-localization (larynx, oral cavity, oropharynx). (**D**) Kaplan-Meier curve stratified after 1^st^ quartile (low), 2^nd^ and 3^rd^ quartiles (medium), and 4^th^ quartile (high) Singscores with Cox proportional hazard ratio (HR), 95% confidence intervals (CI), logrank p-values and survival table is shown for oral cavity cancers of the malignant-basal TCGA patients. (E) Top: Frequencies and percentages of nodal metastases (N0-3) in malignant-basal patients of the TCGA cohort with low (1^st^ quartile), medium (2^nd^ and 3^rd^ quartiles), and high (4^th^ quartile) p-EMT-Singscores including Fisher’s exact test. Bottom: p-EMT-Singscores are plotted against the nodal status ranging from N0 to N3 according to the UICC classification. Included Kendall’s tau and p-value, and Kruskal-Wallis p-value. (**F**) Kaplan-Meier curve stratified after 1^st^ quartile (low), 2^nd^ and 3^rd^ quartiles (medium), and 4^th^ quartile (high) Singscores with Cox proportional hazard ratio (HR), 95% confidence intervals (CI), logrank p-values and survival table is shown for oral cavity cancers of the MD Anderson Cancer Center (MDACC).

Within the malignant-basal subtype of TCGA HNSCC, 90.9% patients (n = 50/55) suffered from oral cavity tumors (**Fig. 2C**). In order to exclude potential confounding drivers within the remaining n = 2 laryngeal and n = 3 oropharyngeal cancer patients, groups of oral cavity cancers only were dichotomized according to p-EMT-Singscores regarding survival analysis. Similarly, p-EMT-Singscores correlated with OS in these patients (High vs. low: HR: 9.17, 95% CI: 1.88 – 44.72, logrank p: 0.005; High vs. medium: HR: 2.89, 95% CI: 1.09 – 7.65, logrank p: 0.057) (**Fig. 2D**).

The incidence of nodal metastases was assessed in patients with low (1^st^ quartile), medium (2^nd^ and 3^rd^ quartiles), and high (4^th^ quartile) p-EMT-Singscores. Fisher’s exact test disclosed a significant inbalance in the proportion of nodal metastases with more incidences of N1, N2, and N3 in patients of medium and high groups compared to low p-EMT-Singscores. A Kendall rank correlation coefficient of tau = 0.298 (p= 0.005) associating the nodal status (N0-3) with increasing p-EMT-Singscores was seen. The p-EMT-Singscore levels between single nodal statuses were significantly different in their mean values (Kruskal-Wallis p = 0.026) (**Fig. 2E**). To further test the value of p-EMT-Singscores as a prognostic factor, an independent cohort of oral cavity cancers (n = 62) from the MD Anderson Cancer Center (MDACC) was analyzed (GEO object GSE42743). Gene expression values from cDNA microarrays were available in combination with the patients’ follow-up information. p-EMT-Singscores were determined in the MDACC oral cavity cohort and followed a normal distribution and dispersion (**Suppl. Fig. 1B**). Calculated p-EMT-Singscores were implemented in a Cox proportional hazard model using the identical stratification strategy as for the TCGA cohort (low= 1^st^, medium= 2^nd^ and 3^rd^, and high= 4^th^ quartiles). OS was displayed in Kaplan-Meier curves, revealing a prognostic value for p-EMT-Singscores in this validation cohort similar to the TCGA malignant-basal patients (High vs. low: HR: 11.30, CI: 2.51 – 50.89, logrank p: 0.001; High vs. medium: HR: 2.36, CI: 1.13 – 4.95, logrank p: 0.025) (**Fig. 2F**).

Hence, p-EMT-Singscores represent a valuable tool to predict OS within malignant-basal HNSCC that are primarily observed in the oral cavity.

### Canonical EMT-TF Slug correlates with p-EMT

In order to address whether p-EMT-Singscores can quantify an incomplete transition of cells to a more mesenchymal state while retaining major epithelial features, the expression of selected epithelial and mesenchymal markers was correlated with p-EMT-Singscore values in TCGA and MDACC patients. In the TCGA malignant-basal patients, Keratin 14, E-cadherin, and Claudin 7 displayed a moderate decline, Rab25 a pronounced decrease, whereas EpCAM showed a minor unexpected increase in patients with higher p-EMT-Singscores (**Figure 3A**, **first panel**). In oral cavity cancer patients of the MDACC cohort, Rab25 and Claudin 7 expression decreased with increasing p-EMT-Singscores while E-cadherin, Claudin7, and EpCAM showed fluctuating expressions (**Figure 3A**, **second panel**). In both cohorts, mesenchymal marker Fibronectin was enhanced with increasing p-EMT-Singscores. Vimentin expression was enhanced with p-EMT-Singscores in the TCGA cohort and was enhanced but show some fluctuation at higher p-EM-Singscore values in the MDACC cohort. The expression of EMT-TFs Slug, Zeb1, and Zeb2 also increased in higher p-EMT-Singscores, with Slug showing comparably higher expression levels than Zeb1 and Zeb2 (**Figure 3A,** third and fourth panels). Hence, p-EMT-Singscores correlated with a retention and partial loss of epithelial markers and an overall gain of mesenchymal markers including the expression of EMT-TFs. With the aim to identify potential regulators of p-EMT in HNSCC, the expression of Slug, Zeb1, and Zeb2 was correlated with the common n = 15 and with the top n = 6 p-EMT genes defined by Puram *et al.* (Puram et al., 2017) in the malignant-basal TCGA and MDACC cohorts. Strongest positive correlation of EMT-TFs with single genes of the common and top p-EMT genes was observed for Slug, whereas Zeb1 and Zeb2 did poorly or not correlate in both cohorts studied (**Fig. 3B**). In the malignant-basal TCGA cohort, a strong correlation of Slug with p-EMT-Singscores was demonstrated in a Spearman’s rank analysis (rho = 0.52; p = 5.9E-05). Zeb1 and Zeb2 correlated less strongly with rho-values of 0.39 (p = 0.0034) and 0.27 (p= 0.045), respectively (**Fig. 3C**). Similar results but even more pronounced differences between the correlations of Slug, Zeb1, and Zeb2 with p-EMT-Singscores were computed in patients of the MDACC cohort. While Slug showed almost identical correlation values (rho = 0.51; p = 2.7E-05), Zeb1 and Zeb2 showed poor correlations (rho = 0.28; p = 0.028 and rho = 0.21; p = 0.11, respectively). Especially on a single gene level compared with the top n = 6 p-EMT genes, negative correlations were seen between Zeb1 and Zeb2 with LAMC2 within the MDACC cohort (**Figure 3D-E**).

**Fig. 3:**
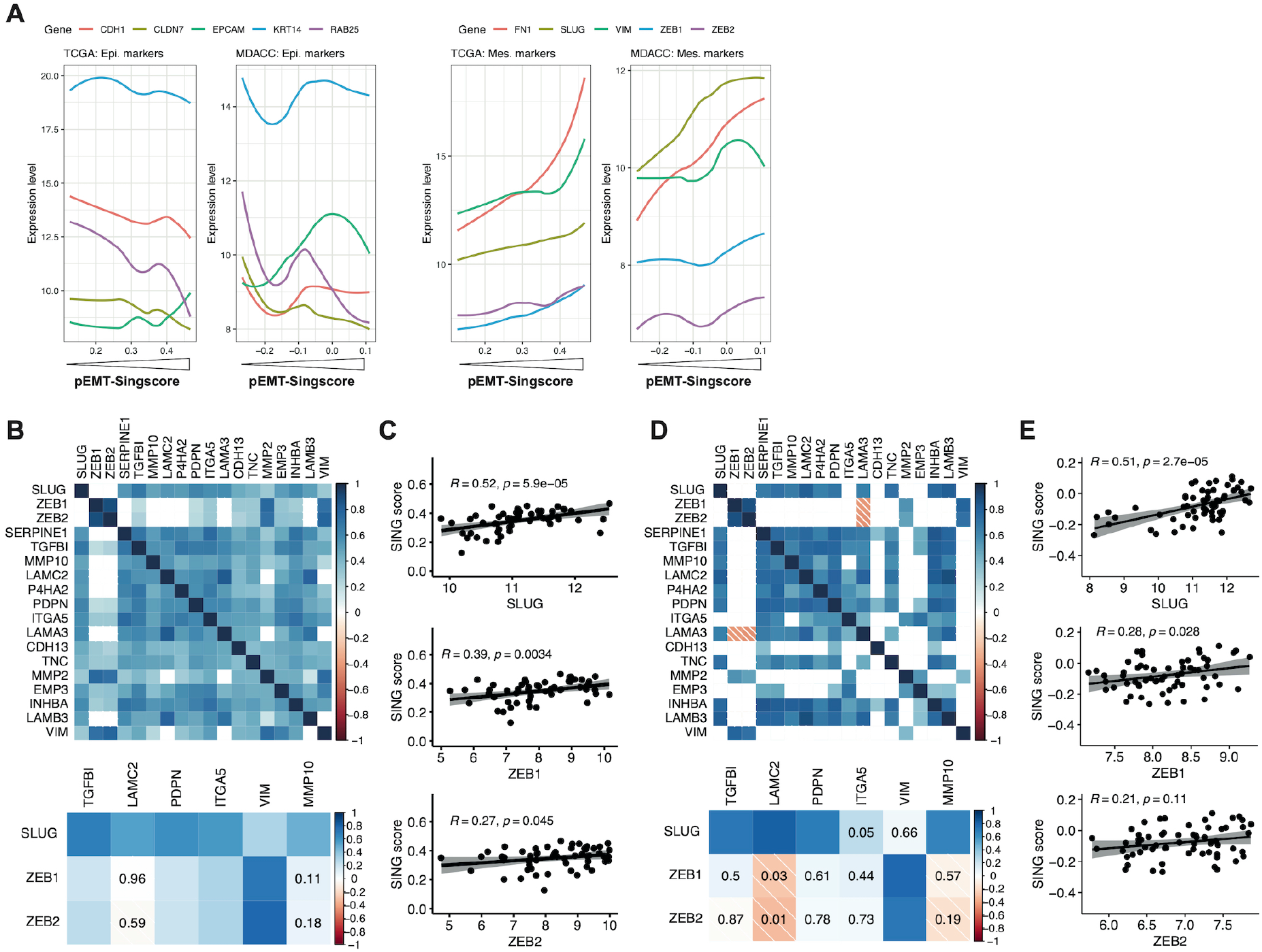
EMT-TF Slug positively correlates with p-EMT-Singscores. (**A**) Expression levels of epithelial (E-cadherin, Claudin 7, Keratin 14, Rab25, EpCAM) and mesenchymal markers (Fibronectin 1, Vimentin), and the EMT-TFs Slug, Zeb1, and Zeb2 in the malignant-basal tumors of the TCGA cohort and the oral cavity cancer MDACC cohort were plotted with line smoothing using loess against p-EMT-Singscores. (**B/D**) Pearson correlation matrices of Slug, Zeb1, and Zeb2 with all n = 15 common p-EMT genes (upper) or the n = 6 top p-EMT genes (bottom) in the malignant-basal patients of the TCGA cohort (B) and oral cavity cancers of the MDACC (E).. Heatmap encodes spearman’s rho. Insignificant values are blank (upper) or indicated (bootom). Significance level ≤ 0.01. (**C/E**) Spearman’s rank correlation of Slug, Zeb1, and Zeb2 with p-EMT-Singscores in the malignant-basal patients of the TCGA cohort (C) and oral cavity cancers of the MDACC (E).

Taken together, the EMT-TF Slug showed a very robust correlation with p-EMT-Singscores and with the common and top p-EMT genes. Therefore, a potential involvement of Slug in the regulation of p-EMT in HNSCC was analyzed *in vitro* in two stable cell lines of the upper aerodigestive tract, *i.e.* FaDu and Kyse30. These cell lines have been chosen on the basis of published results from our own group, demonstrating the induction of EMT through the treatment with high-dose EGF, which was partly controlled *via* activation of Slug transcription (Pan et al., 2018).

### Slug knockout in vitro

Cellular effects of Slug were analyzed *in vitro* following CRISPR/Cas9-mediated knock-out (KO) in FaDu and Kyse30 cells (**Suppl. Fig. 2A**). Wild-type (WT) FaDu and Kyse30 cells, Slug^+/−^ FaDu clone F2.15, Slug^−/−^ clones F2.17 and F2.38, and Slug^−/−^ Kyse30 clone K2.9 were treated with EMT-inducing concentrations of EGF (Pan et al., 2018). Morphologic assessment did not disclose major differences in EMT-related characteristics following EGF treatment (**Suppl. Fig. 2B**). In FaDu cells, no significant differences in epithelial marker E-cadherin were observed following EGF treatment (**Suppl. Fig. 2C**). Snail expression was moderately down-regulated, whereas ZEB1 was comparably up-regulated in WT and Slug-KO FaDu cell lines. Vimentin expression, as an endpoint of EMT, was significantly lower in all Slug-KO clones relative to WT following EGF treatment (**Suppl. Fig. 2C**). Kyse30 Slug-KO cells expressed higher levels of E-cadherin, Snail, and Zeb1, and similar levels of vimentin compared to WT following EGF treatment (**Suppl. Fig. 2C**).

Since literature reports demonstrated severe effects of SLUG-KO *in vivo* on the hematopoietic system and survival in irradiated mice (Inoue et al., 2002; Perez-Losada, Sanchez-Martin, Perez-Caro, Perez-Mancera, & Sanchez-Garcia, 2003), FaDu and Kyse30 SLUG-KO cell lines were irradiated. After irradiation with 2 and 10 Grey (Gy), no differences in alive, dead, and apoptotic cells were observed with an Annexin V/ PI staining 72 hours post irradiation between WT and SLUG-KO clones (Suppl. **Fig. 2D**).

Hence, the totality of the presented findings is in line with a notion that SLUG is dispensable for the induction of p-EMT but may contribute to a full EMT phenotype.

### Slug induces a p-EMT phenotype, invasion, and irradiation resistance

Next, Slug was exogenously over-expressed (Slug-OE) and localized in the nucleus in FaDu and Kyse30 cells (**Fig. 4A,B**). FaDu-Slug-OE cells retained a primarily epithelial phenotype, however with reduced cell-cell junctions, as judged from light microscopy phase contrast micrographs. Kyse30-Slug-OE cells displayed morphological features of p-EMT with reduced cell-cell contact and spindle-shaped morphology (**Fig. 4C**). Loss of cell-cell contact was further corroborated by reduced expression levels of E-cadherin in FaDu-Slug-OE (16.33 % reduction) and Kyse30-Slug-OE cells (and 83.44 % reduction) (**Fig. 4D, E**). FaDu and Kyse30 Slug-OE cells expressed similarly enhanced Slug levels as compared to control cells (55- and 40-fold, respectively). Interestingly, these comparable levels of Slug increase resulted both in significantly higher levels of Zeb1 and vimentin, but the magnitude of Zeb1 and Vimentin increase was one and two orders of magnitude higher in Kyse30 cells, respectively (**Fig. 4F**). Proliferation rates of FaDu-Slug-OE cells remained unchanged, whereas Kyse30-Slug-OE cells displayed a minor proliferation decrease (9.00 % after 48 hours) (**Fig. 4G**).

**Fig. 4:**
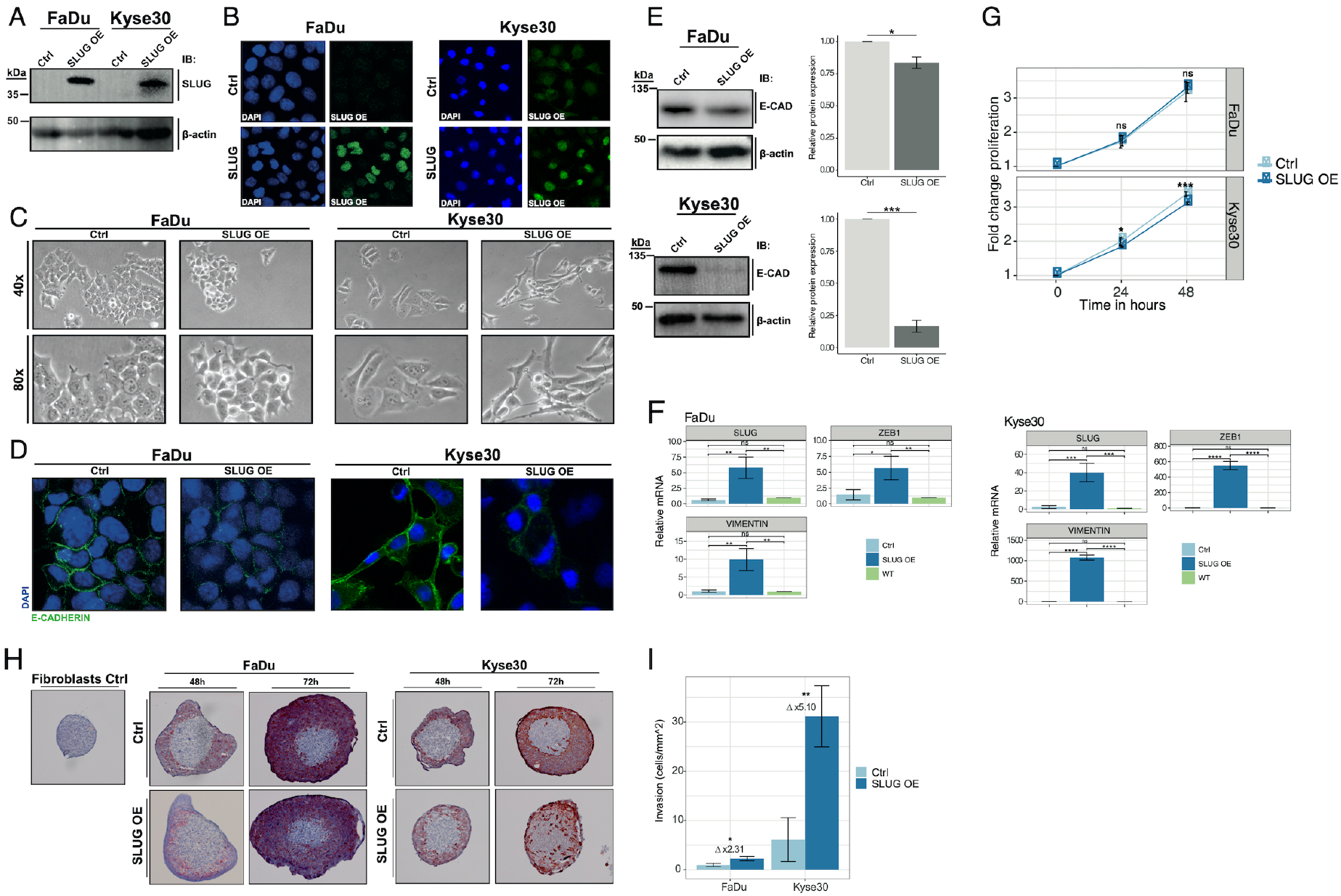
Exogenous SLUG expression induces phenotypic and functional characteristics of p-EMT. **(A)**SLUG was overexpressed (SLUG OE) in FaDu and Kyse30 cells. SLUG expression levels of SLUG OE were assessed and compared to empty vector control cells (Ctrl) by immunoblotting with specific antibodies. (**B**) Expression and localization of SLUG was visualized by immunofluorescence laser scanning microscopy in control (Ctrl) and SLUG OE FaDu and Kyse30 cell lines (green). Nucleic DNA is visualized with DAPI (blue). Shown are representative examples from n = 3 independent experiments. (**C**) Cell morphology after SLUG over-expression was assessed by 40x and 80x magnified light microscopy of vector Ctrl and SLUG OE cells. Shown are representative examples from n = 3 independent experiments. (**D, E**) Expression of E-cadherin (green) in Ctrl and SLUG OE FaDu and Kyse30 cell lines was visualized by immunofluorescence laser scanning microscopy (**D**) and quantified by western blotting of whole cell lysates (**E**, left panel). (**E**, right panel) Shown is a quantification of E-cadherin protein amounts of SLUG OE vs. Ctrl cells from n = 3 independent experiments. Student’s t test. (**F**) mRNA expression levels of SLUG, vimentin, and ZEB1 in Ctrl, SLUG OE, and wildtype (WT) FaDu and Kyse30 cell lines are shown as mean with standard deviations from n = 3 independent experiments performed in triplicates. One-way ANOVA post-hoc Tukey HSD. (**G**) Cell proliferation of Ctrl and SLUG OE FaDu and Kyse30 cell lines over 48 hours are shown as mean from n = 3 independent experiments with n = 6 replicates each. One-way ANOVA post-hoc Tukey HSD. Ns – not significant; * p-value ≤ 0.05; ** p-value < 0.01; *** p-value < 0.001; **** p-value < 0.0001. (**H**) Normal human skin fibroblast spheroids were formed and co-cultured with vector control (Ctrl) or SLUG overexpressing (SLUG OE) FaDu and Kyse30 cell lines. Fibroblast spheroid invasion after 48 and 72 hours was visualized by pan-cytokeratine IHC staining of cryosections. Scale bars are indicated. Shown are representative examples of n = 3 independent experiments with multiple spheroids. (**I**) Invasion was quantified by Matrigel invasion assay. SLUG OE and Ctrl FaDu and Kyse30 cell lines were assessed after 24 hours of invasion by cell counting. Shown are mean and standard deviations of n = 3 independent experiments. Student’s t test.

In the TCGA and MDACC cohort, Slug showed a strong correlation to the top six p-EMT genes vimentin, ITGA5, LAMC2, MMP10, PDPN, and TFGB1 (see **Fig. 3B,D**). Slug over-expression moderately increased the transcription of five from six p-EMT genes in FaDu, namely ITGA5, LAMC2, PDPN, TFGB1, and vimentin (**Suppl. Fig 3**, **Fig. 4F**). In Kyse30 cells, mRNA levels of three out of six genes, namely ITGA5, TGFB1 and vimentin were moderately or strongly increased, respectively. The expression of MMP10 was substantially decreased in Slug-OE cells for unknown reasons (**Suppl. Fig 3**, **Fig. 4F**). Hence, Slug over-expression partially influences the expression of the top p-EMT genes.

Next, the invasive capacity of Slug-OE cells was addressed in a 3D co-culture model in which control- and Slug-OE-FaDu and -Kyse30 cells were added as single cell suspensions to pre-formed human skin fibroblasts spheroids (**Suppl. Fig. 4A**). FaDu and Kyse30 control cell lines (cytokeratin-staining) accumulated around fibroblast spheroids with no obvious signs of invasion (**Fig. 4H**, upper middle and right panel). FaDu-Slug-OE cells showed moderately increased invasion, with single and small aggregates of carcinoma cells infiltrating fibroblast spheroids (**Fig. 4H**, lower middle panel). Kyse30-Slug-OE cells displayed a strong invasive phenotype with numerous invaded cells in the inner area of fibroblast spheroids (**Fig. 4H**, lower right panel).

In a matrigel invasion assay, FaDu-Slug-OE cells showed a 2.31-fold higher invasion compared to controls (2.29 ± 0.45 cell/mm^2^ and 0.99 ± 0.28 cells/mm^2^, respectively) (**Fig. 4I**, **Suppl. Fig. 4B**). Kyse30 cells revealed a 6.19-fold higher invasive potential than FaDu cells. However, Slug-OE fostered matrigel invasion of Kyse30 cells by 5.10-fold compared to Kyse30 control cells (31.17 ± 6.23 cells/ mm^2^ and 6.13 ± 4.45 cells/ mm^2^, respectively) (**Fig. 4I**, **Suppl. Fig. 4B**).

Next, effects of Slug expression on the cells’ sensitivity to apoptosis and irradiation were addressed. Slug expression had no effect *per se* on percentages of alive, dead, and apoptotic cells (**Suppl. Fig. 5A**). Proportions of living cells following induction of apoptosis with 100 nM and 500 nM of Staurosporine were significantly higher in FaDu-Slug-OE cells, whereas the apoptotic fraction was significantly lower compared to control cells (**Suppl. Fig. 5A**). In Kyse30 cells, 100 nM of Staurosporine had no substantial effect on cell apoptosis and survival, however exposure to 500 nM of Staurosporine resulted in significantly higher fraction of living Slug-OE cells to control cells (16.53 % difference). Further, 24.89 % of Kyse30-Slug-OE cells appeared in the dead cell fraction *versus* 45.89 % in the control cell line (**Suppl. Fig. 5A**).

**Fig. 5:**
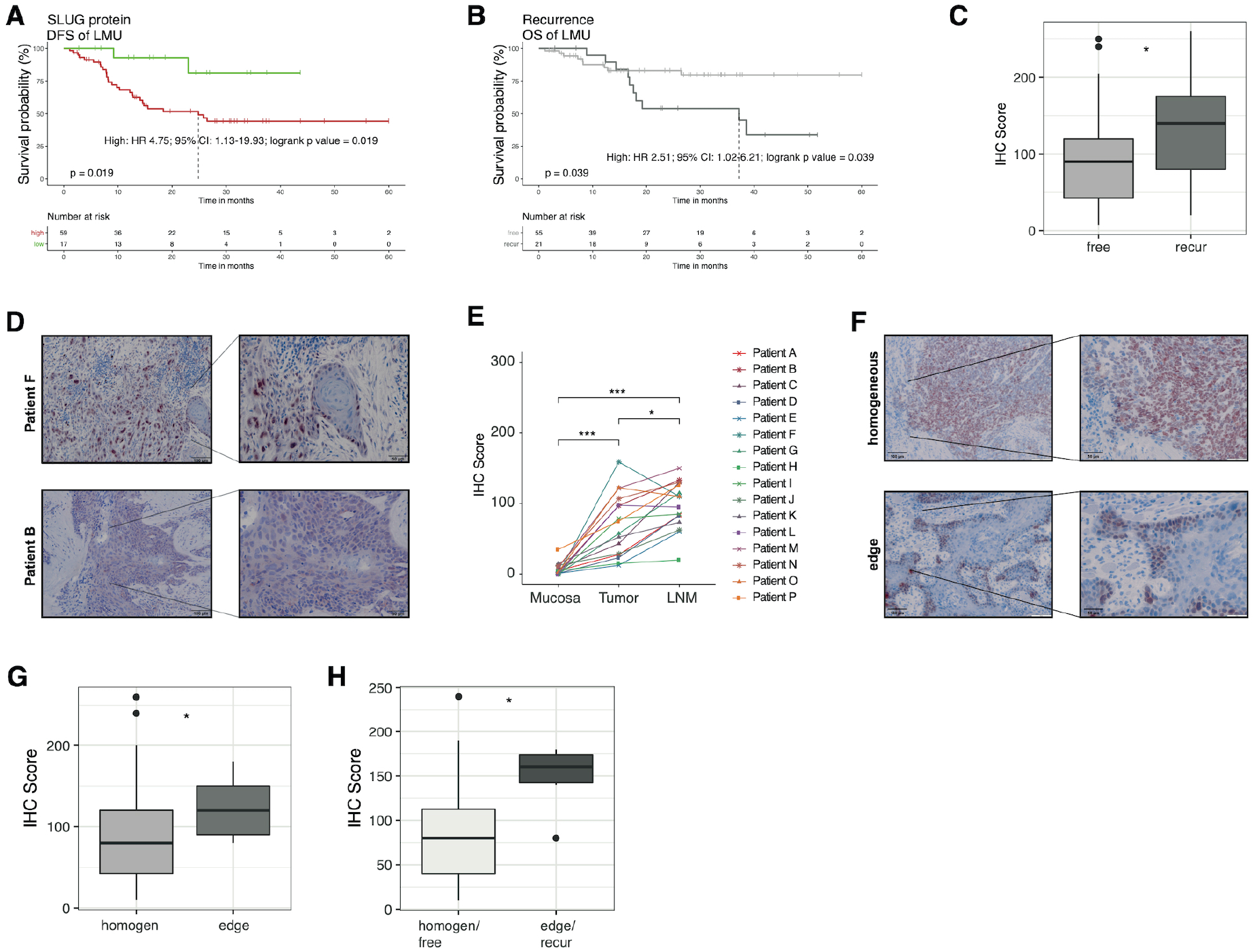
SLUG protein level is prognostic and associated with tumor progression. (**A**) Kaplan-Meier survival curve and tables with logrank p-value, Cox HR, and 95 % CI. LMU cohort of n = 76 HPV-negative HNSCC patients was stratified according to SLUG IHC score. 1^st^ quartile (low) vs. 2^nd^−4^th^ quartiles (high) are shown. (**B**) Kaplan-Meier survival curve and table showing the overall survival based on the occurrence of tumor recurrences. (**C**) Box plots show respective SLUG IHC score of patients suffering from recurrence vs. recurrence-free. Wilcoxon test is shown. * p-value ≤ 0.05. (**D**) Representative examples of high (patient F) and low (patient B) nuclear IHC SLUG staining intensity. Scale bars are indicated. (**E**) Line plots of nuclear SLUG IHC score in tumor-free mucosa, tumor, and lymph-node metastases from n = 16 LMU HNSCC patients. One-way ANOVA post-hoc Tukey HSD is shown. * p-value ≤ 0.05, ** p-value < 0.01, *** p-value < 0.001. (**F**) Representative examples of homogeneous (Patient S) or edge (Patient Q) SLUG staining. Scale bars are indicated. (**G**) Box plots of Slug IHC scores according to protein localization (homogen.: homogeneous or edge). Wilcoxon test is shown. * p-value ≤ 0.05. (**H**) Box plots of Slug IHC scores in patients of the LMU cohort showing a homogeneous Slug expression and no recurrence vs. a Slug expression at the periphery (edge) and recurrences. Wilcoxon test is shown. * p-value ≤ 0.05.

Irradiation of FaDu control and Slug-OE cells with 10 Gy led to significant differences of alive fractions (Ctr.: 62.67 Slug-OE: 80.54 %) and apoptotic fractions (Ctrl.: 27.26 %, Slug-OE: 12.36 %) (**Suppl. Fig. 5B**). Irradiation of FaDu cells with 2 Gy generally did minorly decrease cell survival overall. Significant differences in alive and apoptotic cell fractions between control and Slug-OE cells were observed (alive: 95.55 % vs. 97.13 % / apoptotic: 3.48 % vs. 1.05 %) but were limited (**Suppl. Fig. 5B**). Kyse30 cells were generally more resistant to irradiation than FaDu cells. When irradiated with 10 Gy, 85.29 % of Kyse30 control cells and 92.82 % of Slug-OE cells were contained in the alive fraction, 12.96 % of control cells and 2.80 % of Slug-OE cells were present in the dead fraction (**Suppl. Fig. 5B**). Upon fractionated irradiation of five times 2 Gy, 57.7 % of FaDu control cells were alive and 32.56% dead. In contrast, 72.91 % of FaDu-Slug-OE cells were alive and 19.62 % dead (**Suppl. Fig. 5C**). In Kyse30, fractionated irradiation led to no significant difference between control and Slug-OE cells (**Suppl. Fig. 5C**). Clonogenic assays of irradiated cells confirmed an enhanced resistance of FaDu-Slug-OE cells following 2, 4, and 6 Gy, and enhanced resistance of Kyse30-Slug-OE cells following 4 Gy irradiation (**Suppl. Fig. 5D-E**).

Taken together the data demonstrate that Slug expression promotes p-EMT-associated phenotypic and functional features including enhanced invasion and treatment-resistance.

### Tumor recurrence is associated with Slug expression

Based on the effects of Slug expression on invasion and resistance to therapy *in vitro*, a potential association of Slug protein with recurrence was assessed. For this, protein levels of Slug were quantified in a cohort of HPV-negative primary HNSCC (LMU cohort; n = 76) using immunohistochemical detection and IHC scoring as described (Mack & Gires, 2008). The resulting IHC scores (range 0-300) served to stratify patients. Low Slug protein expression (1^st^ quartile) correlated with significantly longer disease-free survival (DFS) (**Fig. 5A**; HR: 4.75, 95% CI: 1.13-19.93, p = 0.019). Expectedly, patients suffering from any tumor recurrence including local, locoregional, and distant tumors/metastases had a significantly poorer OS (**Fig. 5B**; HR: 2.51, 95% CI: 1.02-6.21, p = 0.039). Concomitantly, patients who suffered from a recurrence expressed significantly higher levels of Slug protein in the index carcinoma compared to patients without any recurring tumor over five years of clinical follow-up (**Fig. 5C**).

### Lymph node metastases show enhanced Slug

A panel of tumor-free mucosa, primary tumor, and lymph node metastases was obtained from n = 16 LMU HNSCC patients and IHC scoring for Slug expression was performed (representative examples in **Fig. 5D)**. Lymph node metastases (IHC score mean: 97.97; median: 102.5) showed significantly enhanced nuclear Slug expression compared to tumor (IHC score mean: 69.92/ median: 66.25) and mucosa (IHC score mean: 6.73/ median: 3.75) (**Fig. 5E**).

EMT at the interface of tumor and stromal areas may facilitate local and distant recurrence through increased migration and invasive (Baumeister et al., 2018; Puram et al., 2017). Slug expression patterns were either homogeneous or enriched at the tumor-stroma interface (edge) (**Fig. 5F**). Edge localization of Slug in tumor samples was associated with significantly higher overall Slug expression compared to tumors with homogeneous Slug expression (**Fig. 5G**). Furthermore, recurrence-free patients with a homogeneous Slug expression were characterized by substantially lower Slug expression than patients with recurrences and a localization of Slug at the edges of tumor areas (**Fig. 5H**).

Hence, a strong and preferably peripheral expression of Slug in primary HNSCC correlated with tumor recurrence, with disease progression, and the formation of lymph node metastases.

## Discussion

HNSCC confront clinicians with severe issues as they are difficult to treat due to a high degree of resistance towards current therapy regimen, frequently yielding metastatic and recurrent disease. Molecular heterogeneity in HNSCC is considered a central driver of therapy resistance influencing clinical outcome (Leemans et al., 2018; Mroz et al., 2015; Mroz et al., 2013; Puram et al., 2017). Partial acquisition of mesenchymal traits emerged as an important mechanism of heterogeneity in oral cavity cancers (Puram et al., 2017) and the results of the present study shed additional light on p-EMT as a common problem in the malignant-basal subtype of HNSCC, i.e. primarily in oral cavity cancers. We have quantified p-EMT using a signature of n = 15 common p-EMT genes defined across patients with oral cavity cancers (Puram et al., 2017) by applying the recently published Single Sample Scoring of Molecular Phenotype (Bhuva et al., 2019; Foroutan et al., 2018), which represents a superior gene set enrichment analysis (GSEA) providing stable Singscores, within the TCGA HNSCC cohort (Cancer Genome Atlas, 2015). Even though not reliant on background samples, Singscores are a GSEA that depends on a sufficiently large number of total genes to quantify enrichment scores for a chosen gene set (n = 15 genes out of a pool of n = 10,000 in the present study). As inferred from scRNAseq of oral cavity cancers, p-EMT is mostly relevant in tumors of the malignant-basal subtype (Puram et al., 2018; Puram et al., 2017). Until today, large-scale scRNAseq screens are not a worldwide standard-of-care for (HNSCC) patients. Thus, in order to adequately transfer the information content of scRNAseq to information from bulk sequencing, we have analyzed HPV-negative patients in the TCGA cohort of the basal and mesenchymal subtypes, only. Using marker genes for non-malignant tumor-resident cells, the influence of non-malignant cells was minimized. The resulting selection of TCGA patients was considered as a surrogate for the malignant-basal subtype. Applying Singscores to the malignant-basal subtype of HNSCC in the TCGA cohort demonstrated that the common p-EMT gene signature, but not multiple random 15 genes sets, correlated with poor OS and with the presence of nodal metastases. In agreement with the Puram study (Puram et al., 2017), malignant-basal HNSCC in the TCGA cohort were in the majority oral cavity tumors. Importantly, results from the TCGA cohort were validated in an independent cohort of oral cavity cancers, demonstrating a general impact of p-EMT on disease progression and clinical outcome in these tumors.

In conclusion, reducing dimensional complexity of p-EMT by Singscoring allowed a distinctive stratification and prognosis of clinical outcome depending on disease sub-groups. Measuring the degree of p-EMT is feasible at the single patient level using Singscores in bulk sequencing information and provides a novel stratification parameter for HNSCC patients in the context of next generation sequencing-based precision medicine (Bhuva et al., 2019). A high p-EMT SING score in patients suffering from oral cavity cancers of the malignant-basal subtype might represent a supportive rationale for full treatment regimens, whereas p-EMT low-risk patients might profit from treatment de-escalation.

Within the group of canonical EMT-TFs, Slug expression correlated best with p-EMT Singscores and common and top p-EMT genes in the TCGA and MDACC cohorts, which is in line with published data on the expression of Slug in p-EMT tumors, although not at the single cell level (Puram et al., 2017). In contrast, Zeb1 and 2 expression was only poorly or not correlated to p-EMT-Singscores and common p-EMT genes. Based on published functions of Slug in EMT regulation (Bolos et al., 2003; Perez-Mancera et al., 2005), we hypothesize that Slug is not only a surrogate for p-EMT but rather actively contributes to induce a p-EMT in HNSCC. This is supported by results showing that Slug expression in cell lines of the head and neck region was sufficient to induce a phenotype and functions, such as invasion and decreased sensitivity to irradiation, that are commonly attributed to p-EMT. A recent meta-analysis of the prognostic value of EMT-TFs in HNSCC disclosed that Twist, Snail, Slug, and Zeb1 correlated with significantly poorer OS (Wan et al., 2020). Further reports demonstrated Snail and Twist as major inducers of EMT in HNSCC (Goppel, Mockelmann, Munscher, Sauter, & Schumacher, 2017; Li et al., 2019). These EMT-TFs were not further investigated in the present study, because they were not expressed to significant levels in the 10,000 common genes of the TCGA cohort. Nonetheless, it cannot be formally excluded that combinatorial functions of Slug, Snail, Twist1, and others orchestrate the complex p-EMT phenotype. However, a meta-analysis across breast cancer studies and a second study across different cancer entities, including HNSCC, named Slug as the main regulator of EMT in patients (Ghulam et al., 2019; Imani, Hosseinifard, Cheng, Wei, & Fu, 2016).

Regulatory mechanisms involved in the induction of Slug expression in HNSCC are poorly understood. Gao *et al.* demonstrated that loss of E-cadherin along with an increased expression of vimentin and Slug at tumor margins was related to the activation of the EGFR/ERK-pathway (Pinilla-Macua, Grassart, Duvvuri, Watkins, & Sorkin, 2017) through the release of EGF by cancer-associated macrophages in the stroma (Gao et al., 2018). In concordance with these findings, our group reported that strong activation of EGFR by EGF resulted in high phosphorylation of ERK and induction of EMT with the up-regulation of Snail, Slug, and Zeb1 (Pan et al., 2018). Accordingly, patients with a high expression of EGFR or of phosphorylated ERK and/or Slug were characterized by poorer OS (Pan et al., 2018). We suggest that over-expression of EGFR, as observed in HNSCC (Kalyankrishna & Grandis, 2006), could promote a strong activation of ERK and the subsequent induction of p-EMT *via* Slug in HNSCC, preferentially at the tumor margin. As a result, p-EMT might confer an increased invasive potential to Slug-positive carcinoma cells allowing them to locally invade and generate deposits of tumor with enhanced therapy resistance that are the seeds of future recurrences and metastases. Both, *in vitro* data from the present study on the enhanced invasive potential of Slug-positive carcinoma cells and improved resistance to irradiation, and the positive correlation of Slug expression *in vivo* with recurrences, in particular local recurrences, are in support of this notion.

In summary, p-EMT is a main parameter of tumor progression that can be quantified using Singscores and that strongly correlates with Slug expression in HNSCC. The development of prognostic p-EMT signatures based on Singscores might be applied in the clinical setting to stratify patients more precisely into subgroups with differing risk of disease spread and recurrence. Patients at high risk would profit from a longitudinal monitoring and could, potentially, benefit from more aggressive treatments to suppress tumor recurrence.

## Materials and Methods

### Data analysis

Data analysis was performed using R (R Core Team, R: A Language and Environment for Statistical Computing, R Foundation for Statistical Computing, 2017; R version 3.6.1 (2019-07-05)). Correlation matrices were calculated and illustrated with CRAN *corrplot* package. Univariate survival analysis was performed with CRAN packages *survival* and *survminer*. Unless otherwise stated, further analysis was performed with built-in packages and functions from the CRAN package *tidyverse.*

### TCGA patient selection

Top 10,000 protein coding genes from the whole human genome across all TCGA patients were defined (Cancer Genome Atlas, 2015). The R-package *CGDS* (Bioconductor) was used to extract expression profiles (cancer study: “hnsc_tcga_pub”/ case list: “hnsc_tcga_pub_all”/ genetic profile: “hnsc_tcga_pub_rna_seq_v2_mrna”). A total of 10,000 complete mRNA profiles could be extracted for n = 243 HPV-negative TCGA HNSCC patients. Molecular subtypes for each TCGA patient were extracted and patient data was processed in accordance to Puram et *al.* (Puram et al., 2017). Briefly, expression values of the extracted top 10,000 protein coding genes across all patients were log2 transformed, centered, and a correlation analysis across all patients was computed. Patients with a mean Pearson correlation > 0.1 within their respective subtype and < 0.1 compared to all patients from the other subtypes were kept for further analysis. Then resulting n = 46 patients from the mesenchymal and n = 38 from the basal-like subtype group were applied to filter out patients with highest influence of non-malignant cells. Therefore, centered gene expression values from indicated marker genes was applied to compute an Euclidean distance-to-distance matrix using the *dist* function from the R CRAN *stats* package. Then hierarchical clustering was applied using the *hclust* function from *stats* package. The resulting clustering tree was cut at a height of 13 resulting in 4 different clusters. The cluster with the largest influence of non-malignant cells relative across all patients consisting of n = 29 mesenchymal tumors was removed. All others were kept for further analysis (n = 55).

### Computing Singscores of TCGA and MDACC patients

As aforementioned, gene expressions of 10,000 protein coding genes across all TCGA patients were extracted (Cancer Genome Atlas, 2015). A total of 10,000 complete mRNA profiles could be extracted for n = 243 HPV-negative TCGA HNSCC patients. Expression data was Log2 transformed prior to calculations. Using the R-package *singscore* (Foroutan et al., 2018) (Bioconductor), a Singscore was computed for each TCGA patient with the common p-EMT genes defined by Puram et *al*. (n = 15 genes) (Puram et al., 2017). For univariate survival analysis, Cox-proportional hazard ratios (HR) > 1 with logrank p-value ≤ 0.05 was accepted as relevant. To test the validity of results obtained with the p-EMT gene set, n = 10,000 random sets of 15 genes from the extracted gene pool (n= 10,000, excluding all n = 100 p-EMT genes) served to compute Cox-proportional hazard models with log-rank p-values. For visualization, Singscores were implemented to dichotomize patients into 25 % lowest (“low”, 1^st^ quartile), intermediate 50 % (“medium”, 2^nd^ and 3^rd^ quartiles), and 25 % highest groups (“high”, 4^th^ quartile). Then, a Cox-proportional hazard model, median survival times, and log-rank p-values were calculated and included in Kaplan-Meier plots. MDACC data was retrieved from the GEO object GSE42743 by using the R Bioconductor packages *GEOquery, affy, AnnotationDbi, and hgu133plus2.db* to extract and map the microarray expression data, and to receive the according clinical data set. The *affy* function *rma* for robust multi-array average expression measure with default settings was applied to MDACC transcriptome data to receive log2 transformed data. From the 10,000 protein coding genes, n= 9831 were recovered in the MDACC data set. Then, as afore described for the TCGA data, a Singscore with n= 15 common pEMT was computed, tested in a Cox-propotional hazard model, applied to patient dichotomization, and visualized by a Kaplan-Meier including a Cox-proportional hazard model, median survival times, and log-rank p-values.

### Human samples and ethics statement

The Ludwig-Maximilians-Universität (LMU) of Munich, Germany, HNSCC cohort included HPV-negative tumor specimens from n = 76 patients (Pan et al., 2018). Clinical samples were obtained after written informed consent during routine surgery, based on the approval by the ethics committee of the local medical faculties (Ethikkommission der Medizinischen Fakultät der LMU; #087–03; #197–11; #426–11) and in compliance with the WMA Declaration of Helsinki and the Department of Health and Human Services Belmont Report. Samples were cryo-preserved by snap-freezing in tissue-Tek^®^ (Sakura, Finetek, The Netherlands) and processed to 5 μm thick sections for further staining as has been described in (Baumeister et al., 2018).

### Immunohistochemistry scoring and immunofluorescence

Specific antibodies against Slug (C19G7, Cell Signaling Technology, NEB, Frankfurt, Germany, #9585, 1:400), pan-cytokeratine (polyclonal, Invitrogen, Camarillo, USA, #18-0059, 1:200) and E-cadherin (24E10, Cell Signaling Technology, NEB, Frankfurt, Germany, #3195, 1:400) were used for immunohistochemistry (IHC) and immunofluorescence staining in combination with the avidin-biotin-peroxidase method (Vectastain, Vector laboratories, Burlingame, CA, US) or Alexa Fluor-488-conjugated secondary antibody. Confocal microscopy images were recorded with a TCS-SP5 system (Leica Microsystems; Wetzlar, Germany). IHC scores were formed following staining of cryosections (5 μm) of HNSCC samples by a two-parameters system, which implemented scoring of all specimen in percentages of cells and their staining intensities from 0 to 3 (0= negative, 1= mild intensity, 2= moderate intensity, 3= strong intensity, score = sum(% x intensity); resulting max. score 300) as described (Mack & Gires, 2008). At least two experienced scorers evaluated IHC specimen independently and in blinded manner.

### Cell lines and treatments

FaDu and Kyse30 cell lines were obtained from ATCC and DSMZ and were confirmed by STR typing (Helmholtz Center, Munich, Germany). For SLUG knockout, CRISPR-Cas constructs U6-gRNA:CMV-eCas9-2a-tRFP (p05 transfection plasmid, Sigma-Aldrich) targeting human SLUG exon2 (target sequence: -CGGTAGTCCACACAGTGAT-) were used. Transfected cells were sorted for expression of turbo red fluorescence protein (tRFP) using fluorescence-activated cell sorting (FACS). The bulk of one percent highest tRFP expressing cells was plated into 96-well plate at 1 cell/ well. The knockout status of selected single cell clones was assessed by DNA sequencing and Western blotting for SLUG protein. For SLUG over-expression, Kyse30 and FaDu cells were stably transfected with SLUG-Myc in the 141-pCAG-3SIP vector with MATra transfection reagent (PromoCell) using 1 μg/ mL puromycin (Sigma) for selection. Control cell lines were transfected with an empty 141-pCAG-3SIP vector. Wildtype and knockout cells were maintained in RPMI 1640 or DMEM, 10 % FCS, 1 % penicillin/streptomycin, in a 5 % CO2 atmosphere at 37 °C, and 1 μg/mL puromycin for SLUG over-expressions and pCAG controls. Treatment with 50 ng/mL EGF (PromoCell PromoKine, Heidelberg, Germany) was conducted in respective FCS-free medium after serum starvation over-night.

### Western blotting

Whole cell lysates were extracted with phosphate-buffered saline (PBS) containing 2 % Triton X-100 and protease inhibitors (Roche Complete, Roche Diagnostics, Mannheim, Germany). Protein concentrations were determined by BCA-assay (Thermo Scientific, Schwerte, Germany). Ten to 50 μg of proteins were separated by 10 % SDS-PAGE and visualized with primary Slug or E-cadherin antibody (C19G7, Cell Signaling Technology, #9585, 1:1000, overnight at 4°C / 24E10, Cell Signaling Technology, #3195, 1:1000, overnight at 4°C) and horseradish peroxidase (HRP)-conjugated secondary antibodies (1:5000, 1 hour at room temperature), and the ECL reagent (Millipore, Darmstadt, Germany) in a Chemidoc XRS imaging system (Bio-Rad, Munich, Germany). An HRP-conjugated antibody was used to visualize beta-actin (sc-47778 HRP, Santa Cruz).

### Reverse transcription qPCR analysis

Total RNA was extracted using RNeasy Mini kit (Qiagen) and reverse transcribed with QuantiTect Reverse Transcription kit (Qiagen). The resulting cDNAs were used for analysis with SYBR-Green master mix and LightCycler480 (Roche) or QuantStudio3 (ThermoFisher). Quantifications exceeding a cycle threshold (CT) of 35 were regarded as not expressed. All quantified values were normalized to internal GAPDH control. The relative expression value for each target gene compared to the calibrator for that target was calculated as 2^-(Ct-Cc)^ as described by Livak et *al. (Livak & Schmittgen, 2001)* (Ct and Cc are the mean threshold cycle differences after normalizing to GAPDH).

### Primers used for qPCR quantification

*E-CADHERIN-FW 5’-TGC CCA GAA AAT GAA AAA GG-3’*

*E-CADHERIN-BW 5‘-GTG TAT GTG GCA ATG CGT TC-3’*

*GAPDH-FW 5’-AGG TCG GAG TCA ACG GAT TT-3’*

*GAPDH-BW 5’-TAG TTG AGG TCA ATG AAG GG-3’*

*ITGA5-FW 5’-GGCTTCAACTTAGACGCGGAG-3’*

*ITGA5-BW 5’- TGGCTGGTATTAGCCTTGGGT-3’*

*LAMC2-FW 5’- CAAAGGTTCTCTTAGTGCTCGAT-3’*

*LAMC2-BW 5’- CACTTGGAGTCTAGCAGTCTCT-3’*

*MMP10-FW 5’- TCAGTCTCTCTACGGACCTCC-3’*

*MMP10-BW 5’- CAGTGGGATCTTCGCCAAAAATA-3’*

*PDPN-FW 5’- ACCAGTCACTCCACGGAGAAA-3’*

*PDPN-BW 5’- GGTCACTGTTGACAAACCATCT-3’*

*TGFB1-FW 5’- CTTCGCCCCTAGCAACGAG-3’*

*TGFB1-BW 5’- TGAGGGTCATGCCGTGTTTC-3’*

*SLUG-FW 5‘-TGA TGA AGA GGA AAG ACT ACAG-3’*

*SLUG-BW 5’-GCT CAC ATA TTC CTT GTC ACA G-3‘*

*SNAIL-FW 5‘-GCG AGC TGC AGG ACT CTA AT-3‘*

*SNAIL-BW 5’-CCT CAT CTG ACA GGG AGG TC-3‘*

*VIMENTIN-FW 5‘-GAG AAC TTT GCC GTT GAA GC-3‘*

*VIMENTIN-BW 5’-GCT TCC TGT AGG TGG CAA TC-3‘*

*ZEB1-FW 5’-TGC ACT GAG TGT GGA AAA GC-3’*

*ZEB1-BW 5’-TGG TGA TGC TGA AAG AGA CG-3’*

### Annexin V-FITC and propidium iodide dual staining

To determine effects of staurosporine treatment (dissolved in DMSO) or irradiation, the annexin V-FITC and propidium iodide (PI) dual staining kit (Invitrogen, eBioscience Annexin V-FITC Apop, BMS500FI-300) was used according to manufactures protocol. Briefly, 1×10^6^ harvested cells were resuspended in 1x binding buffer. Five μL annexin V-FITC in 200 μL 1x binding buffer were added and incubated in the dark for 10 minutes. Cells were washed with binding buffer and resuspended in 200 μL 1x binding buffer containing 10 μL PI. Then, 200 μL of 1x binding buffer were added and suspension was analyzed by flow cytometry (Beckman-Coulter, CytoFLEX). Flow cytometry gates were chosen based on untreated controls for each cell line and values were normalized to controls.

### Cell proliferation assay

Cells were counted using a Leica DMi8 microscope with LAS X software and FIJI. In a 96-well, 2,000 cells were seeded initially for each timepoint. Cells were left overnight to fully attach to the plate. The next day (timepoint 0 hours), and 24 or 48 hours later (timepoints 24 and 48 hours) were measured. For counting, cells were stained with Hoechst 33342 dye (ThermoFisher) for 15 minutes. Then, using the LAS X software, 72 images per well with 100× magnification were taken and merged. In FIJI, images in greyscale (16-bit) were compressed to 8-bit by threshold adjustment (in FIJI: Image>Adjust>Threshold>Apply) to remove noise. Then, Watershed function was applied to cut any artificially merged pixels (in FIJI: Process>Binary>Watershed). Finally, resulting particles, representing single cells, were counted and summarized (in FIJI: Analyze>Analyze Particles, setting Size:0-Infinity, Circularity: 0-1, tick “Clear results” and “Summarize”). Resulting counted cell numbers were then analyzed using RsSoftware.

### Fibroblast spheroid invasion assay

Spheroids of normal human foreskin fibroblasts (PromoCell, C-12352) were grown in Ultra Low Attachment plates (ULA) over 24 hours by seeding 1×10^4^ cells in standard DMEM. Following the formation of fibroblast spheroids, 1×10^4^ FaDu or Kyse cells were added and co-cultured for additional 48 and 72 hours. Co-cultured spheroids were carefully harvested with a cut 100 μL pipette tip and immediately frozen in tissue-TEK (Sakura Europe) in a cryomold with liquid nitrogen. Then, cryosections of 5 μm thickness were generated and IHC staining was conducted.

### Matrigel invasion assay

Matrigel invasion assays were conducted in accordance to Shaw *et al.* (Shaw, 2005). Briefly, a total of 1×10^5^ of tumor cells was seeded in 1:10 diluted matrigel-coated 24-well membrane chambers (Corning, cell culture inserts, 8 μm pore size, 353097) in serum-free medium. 24-well chambers contained standard medium. After 24 hours of invasion, cells attached on the top were swiped off with a cotton swab and membranes were carefully extracted with a scalpel.

Then, cells were fixed with methanol and membranes stained with crystal violet and invaded cells were counted visually.

### Clonogenic survival assay

For FaDu, 1×10^3^ and for Kyse30 5×10^3^ cells were plated on a 6-well plate and irradiated the next day. After 14 days for FaDu and 10 days for Kyse30, cells were fixed and stained with crystal violet solution containing methanol. The whole 6-well plate was photographed using the Chemidoc XRS imaging system. Then, to quantify the area of colonies, the ColonyArea Image J Plugin by Guzmán et *al.* (Guzman, Bagga, Kaur, Westermarck, & Abankwa, 2014) was used. Clonogenic survival was calculated by measuring the area of colonies of irradiated relative to non-irradiated control plates.

## Supporting information

Supplementary Figures Schinke et al.

## List of abbreviations

scRNAseq: single cell RNA sequencing
HNSCC: Head and neck squamous cell carcinoma
p-EMT: partial epithelial-to-mesenchymal transition
HPV: Human Papillomavirus
OS: Overall survival
DFS: Disease-free survival
EMT-TF: EMT-transcription factor
ZEB1/2: Zinc finger E-box-binding homeobox 1 and 2
TCGA: The cancer genome atlas program
MDACC: MD Anderson cancer cohort
HR: Hazard ratio
CI: Confidence intervall
LMU: Ludwig-Maximilians University
IHC: Immunohistochemistry
qPCR: quantitative polymerase chain reaction
CRISPR: Clustered regularly interspaced short palindromic repeats
Slug-OE: Slug-overexpression
Ctr: Control
ITGA5: Integrin alpha 5
LAMC2: Laminin subunit gamma 2
PDPN: Podoplanin
TGFB1: Transforming growth factor beta 1
GSEA: Gene set enrichment analysis

## Declarations

### Ethics approval and consent to participate

Clinical samples of the LMU HNSCC cohort were obtained after written informed consent during routine surgery, based on the approval by the ethics committee of the local medical faculties (Ethikkommission der Medizinischen Fakultät der LMU; 087–03; 197–11; 426–11, EA 448-13, and 17-116) and in compliance with the WMA Declaration of Helsinki and the Department of Health and Human Services Belmont Report.

### Consent for publication

Not applicable.

### Data availability

The datasets generated during and/or analyzed during the current study are either publicly available (TCGA (Cancer Genome Atlas, 2015), MDACC GEO object GSE42743) or are available from the corresponding author on reasonable request.

All codes and R-packages used in the study are publicly available and have been disclosed in the Materials and Methods section or are available from the corresponding author on reasonable request.

### Competing interests

The authors declare no conflict of interest. The funders had no role in the design of the study; in the collection, analyses, or interpretation of data; in the writing of the manuscript, or in the decision to publish the results.

### Funding

This research was partly funded by the German Research Council (Deutsche Forschungsgemeinschaft), grant number Gi 540-3/2 and the Bayerisches Staatsministerium für Wirtschaft, Energie und Technologie BayBIO (BIO1803-0003), and the program “Förderprogramm für Forschung und Lehre” of the Ludwig-Maximilians University of Munich (Reg.Nr. 1019/995) to FS.

### Author Contributions

Conceptualization, HS and OG; Methodology, OG, HS and PB; Software, HS; Validation, HS, OG and PB; Formal analysis, HS, MA, MP, JZ; Investigation, HS, MA, OG; Resources, GK, DL, PB, OG, MC, FS, and HS; Data curation, HS, MA, JZ, FS and PB; Writing original draft preparation, HS; writing review and editing, OG, MC and PB; Visualization, HS, OG and MA; Supervision, OG and PB; Project administration, OG and PB; Funding acquisition, OG and FS.

## Acknowledgments

We thank Kristian Unger and Daniel Samaga (Helmhotz Center Munich) for their support and consultation with data analysis in *R*.

## Notes

### Competing Interest Statement

The authors have declared no competing interest.

