## Supplementary Figures Schinke et al. for "Partial epithelial-to-mesenchymal transition is prognostic and associates with Slug in head and neck cancer"

### ABSTRACT

Therapy resistance leading to local recurrence and metastases remains highly problematic in head and neck squamous cell carcinomas (HNSCC). Single cell RNA-sequencing defined a partial epithelial-to-mesenchymal transition (p-EMT) signature associated with metastases in HNSCC. However, the prognostic value of the p-EMT signature and potential drivers of p-EMT in HNSCC remain unclear. Here, single sample scoring of molecular phenotypes (Singscoring) served to establish clinical p-EMT-Singscores that were significantly associated with nodal metastases and predicted overall survival in two independent HNSCC cohorts. p-EMT-Singscores correlated most strongly with EMT transcription factor (EMT-TF) Slug. *In vitro*, Slug promoted p-EMT, enhanced invasion, and resistance to irradiation. In patients, Slug protein levels in tumors predicted disease-free survival and its peripheral expression at the interphase to tumor-microenvironment was significantly increased in recurring patients. Thus, p-EMT represents a novel clinical risk-predictor that impacts on HNSCC patients´ outcome and is partly controlled by Slug.

### Supplementary Information

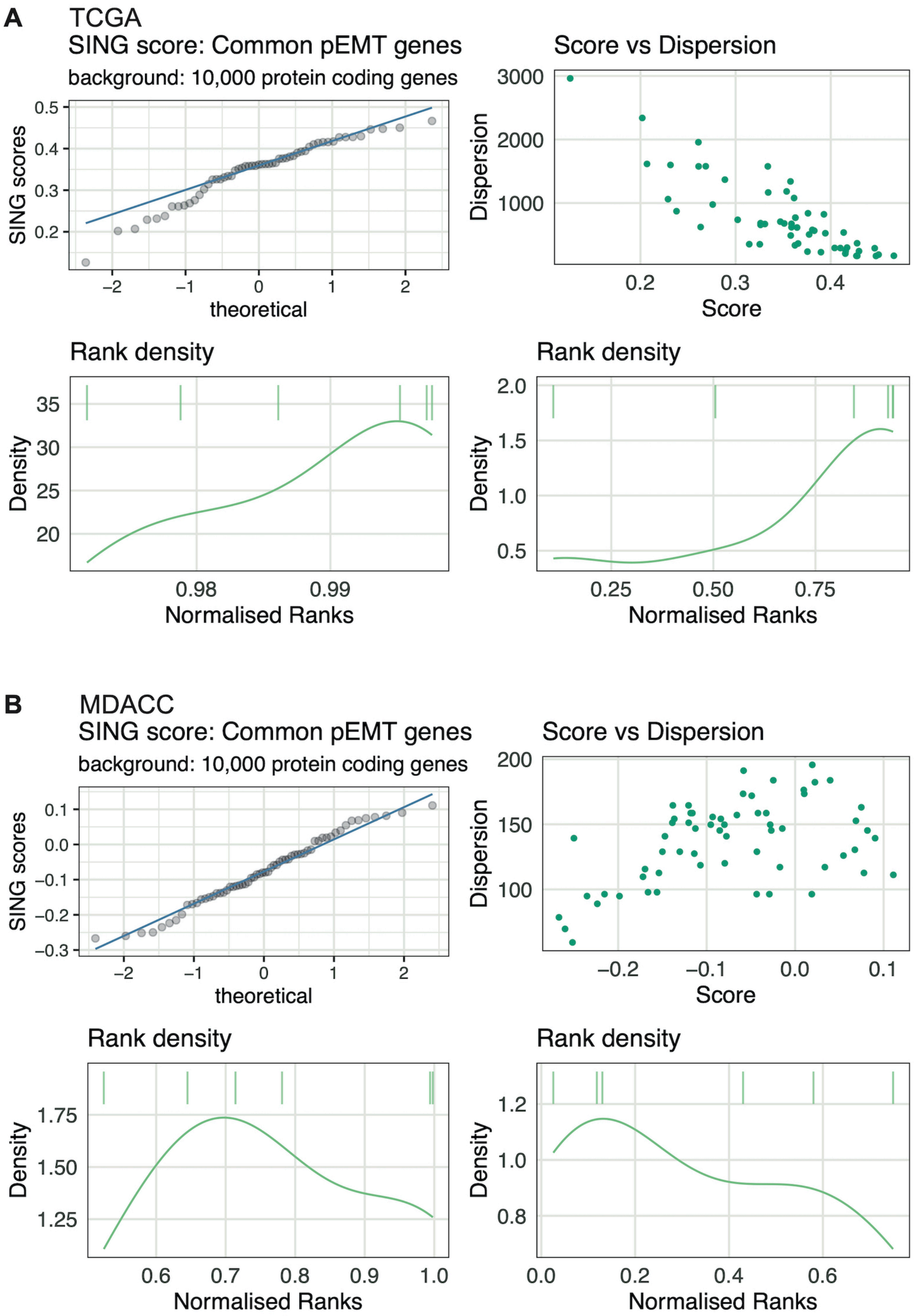


**Suppl. Fig. 1: Computation of p-EMT-Singscores based on n = 15 common p-EMT genes in TCGA cohort.** (**A**) **Upper left**: Quantile-quantile plot of p-EMT-Singscores of common p-EMT genes vs. theoretical quantiles within the TCGA HNSCC cohort. **Lower**: Rank density plot of p-EMT-Singscores from common p-EMTgenes shows patients with lowest (right) and patient with highest p-EMT-Singscore (left) in normalized gene ranks. **Upper right**: Dotplot of dispersion against common p-EMT-Singscores of each TCGA patient. (**B**) Same as (**A**) for patients of the MDACC oral cavity cancer cohort.

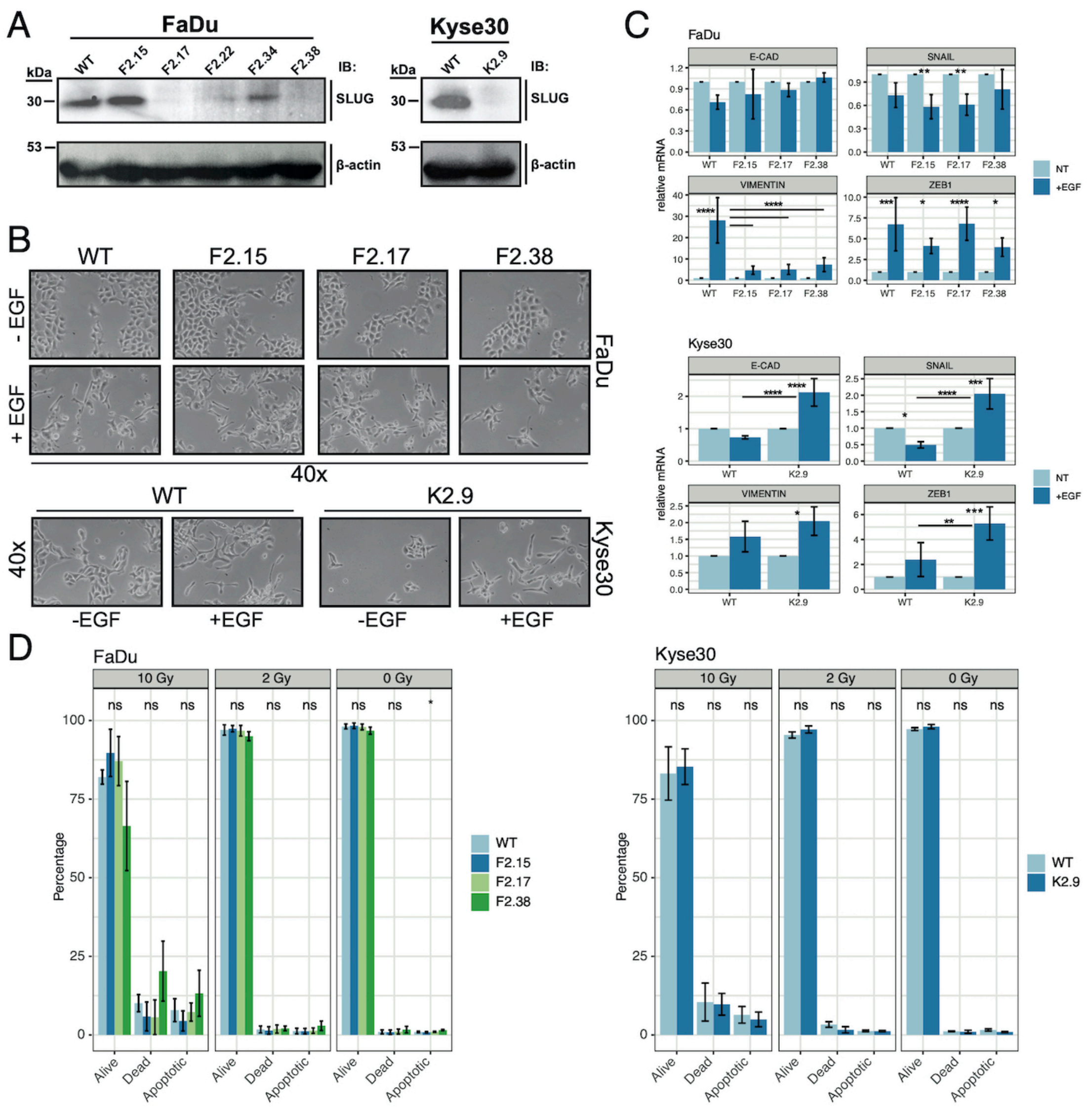


**Suppl. Fig. 2: SLUG knockout in head and neck cell lines.** (**A**) Western blot of wildtype (WT) and SLUG knockout (KO) FaDu and Kyse30 cell lines treated with EGF 50 ng/mL for 72 hours. SLUG expression was assessed by immunoblotting. (**B**) WT and KO FaDu and Kyse30 cells were treated with or without 50 ng/mL EGF. Cell morphology was assessed after 72 hours. Shown are representative micrographs at 40x magnification from n = 3 independent experiments. (**C**) qRT-PCR quantification of the indicated mRNA species in WT and KO FaDu and Kyse30 cell lines treated with 50 ng/mL EGF for 48 hours (normalized to untreated control of each cell line). E-cadherin (E-CAD), SNAIL, vimentin, and ZEB1 mRNA levels were measured. One-way ANOVA post-hoc Tukey HSD as indicated. (**D**) Flow cytometry quantification of Annexin V/ PI staining after 10, 2 and 0 Grey (Gy) irradiation of WT and KO FaDu and Kyse30 cell lines. One-way ANOVA post-hoc Tukey HSD. Shown are results from at least n = 3 independent experiments. Ns – not significant; * p-value ≤ 0.05; ** p-value < 0.01; *** p-value < 0.001; **** p-value < 0.0001.

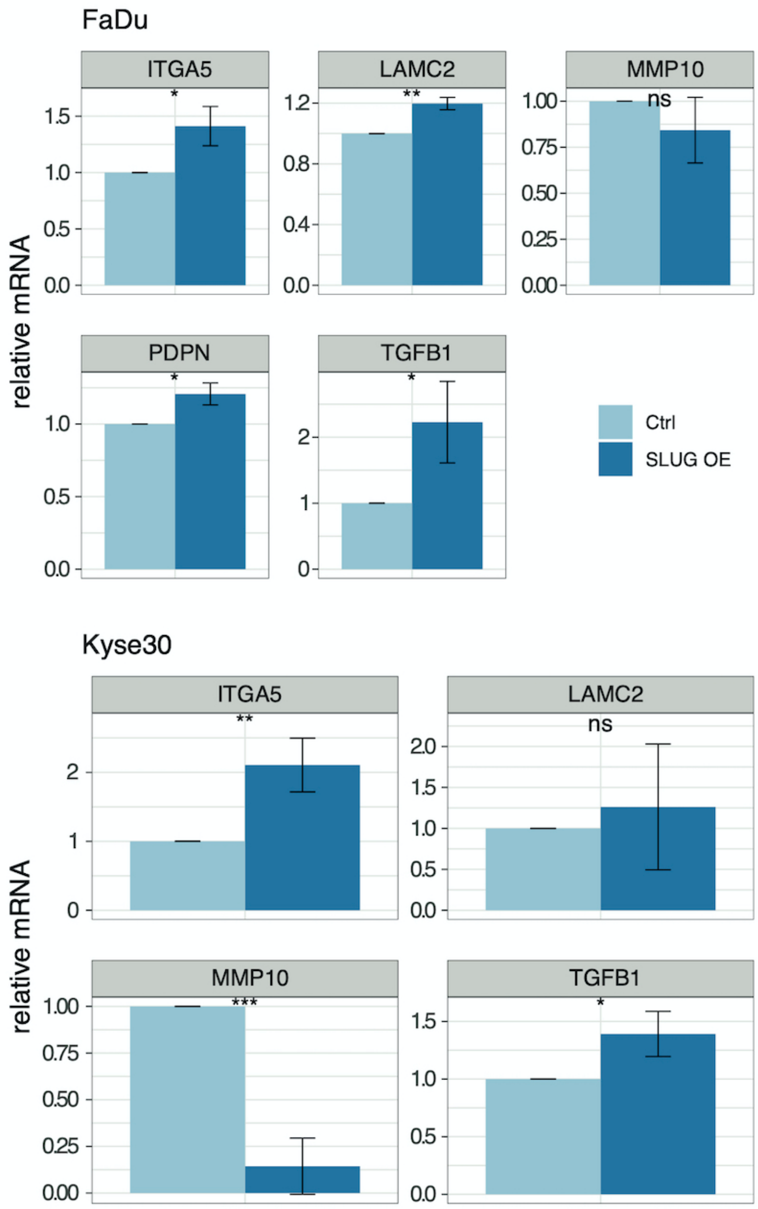


**Suppl. Fig. 3: Top p-EMT genes and SLUG over-expression.** RT-qPCR mRNA quantification of top p-EMT genes ITGA5, LAMC2, MMP10, PDPN, and TGFB1 of vector control (Ctrl) and SLUG overexpressing cells (SLUG OE) FaDu and Kyse30 cell lines. Normalized to vector control cells (Ctrl). Student's t test. Ns – not significant; * p-value ≤ 0.05; ** p-value < 0.01; ***.

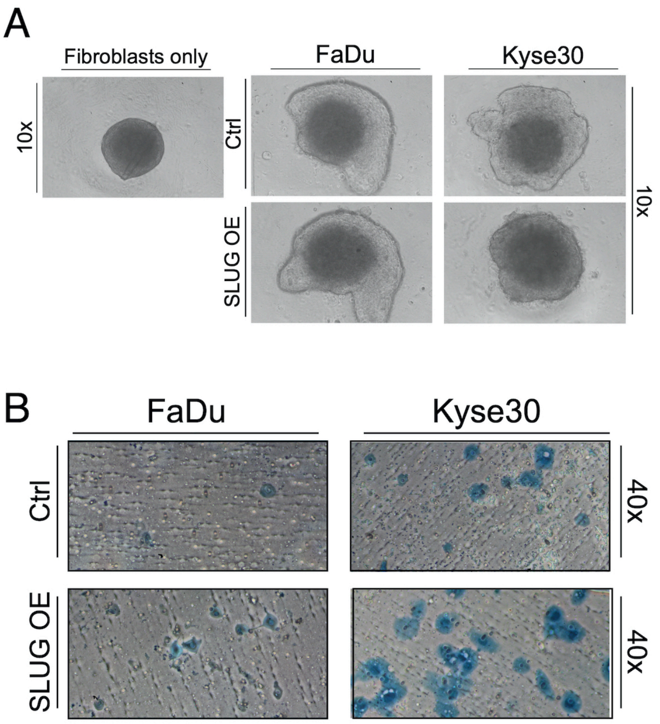


**Suppl. Fig. 4:** (**A**) Light microscopic images with 10x magnification of normal human skin fibroblasts spheroids (left) and co-cultured SLUG overexpressing (SLUG OE) and vector control (Ctrl) FaDu and Kyse30 cells (middle and right) after 72 hours. (**B**) Light microscopic images with 40x magnification of SLUG OE and Ctrl FaDu and Kyse30 cells invaded onto the bottom of membranes from Matrigel invasion assays after 24 hours. Cells were fixed and stained with crystal violet. Shown are representative images of n = 3 independent experiments.

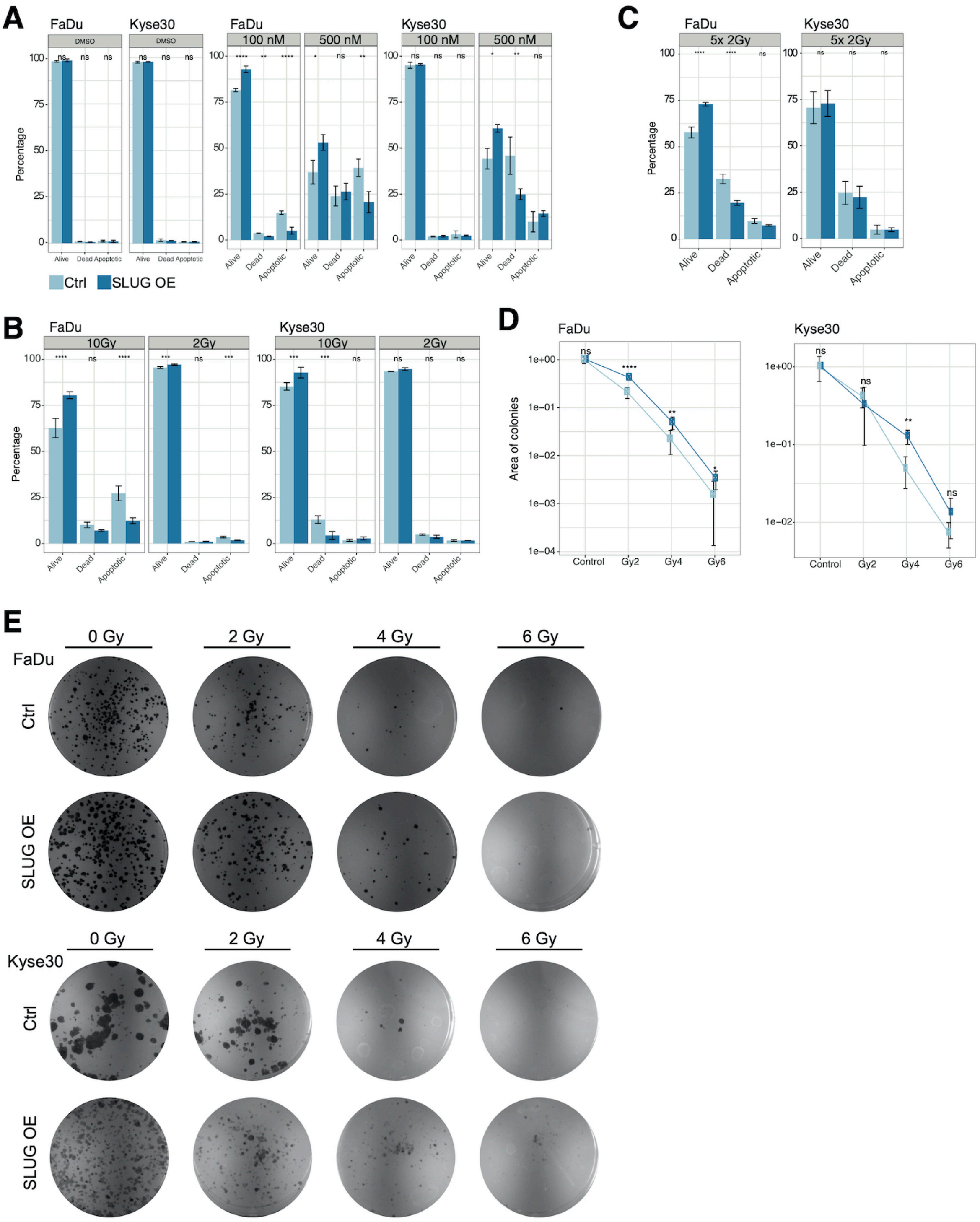


**Suppl. Fig. 5:** (**A**) SLUG OE and Ctrl FaDu and Kyse30 cellline s were treated for 24 hours with 100 and 500 nM of Staurosporine. Cell death was assessed by flow cytometry and Annexin V/ PI staining. Shown are mean and standard deviations of n = 3 independent experiments. (**B-C**) SLUG OE and Ctrl FaDu and Kyse30 cell lines were irradiated with 10 and 2 Grey (Gy) and after 72 hours cell death was assessed by Annexin V/ PI staining. 10 Gy were also applied as fractionation in 5x 2Gy shown in (**C**). Shown are mean and standard deviations of n = 3 independent experiments. (**D**) Clonogenic survival assay of SLUG OE vs. Ctrl FaDu and Kyse30 cell lines with 0 (Control), 2, 4, and 6 Gy irradiation. Area of colonies was measured by ColonyArea ImageJ Plugin after 2 weeks for FaDu and 10 days for Kyse30 cells. Shown are mean and standard deviations of n = 3 independent experiments. (C-F) one-way ANOVA post-hoc Tukey HSD. Ns – not significant; * p-value ≤ 0.05; ** p-value < 0.01; ***. (**E**) BW images of 6-well plates after 14 days (FaDu) or 10 days (Kyse30) of SLUG OE and Ctrl cells with different doses of irradiation as stated. Shown are representative results from n= 3 independent experiments.
